# Only cortical prediction error signals are involved in visual learning, despite availability of subcortical prediction error signals

**DOI:** 10.1101/2023.11.13.566726

**Authors:** Dongho Kim, Zhiyan Wang, Masamichi Sakagami, Yuka Sasaki, Takeo Watanabe

## Abstract

Both the midbrain systems, encompassing the ventral striatum (VS), and the cortical systems, including the dorsal anterior cingulate cortex (dACC), play roles in reinforcing and enhancing learning. However, the specific contributions of signals from these regions in learning remains unclear. To investigate this, we examined how VS and dACC are involved in visual perceptual learning (VPL) through an orientation discrimination task. In the primary experiment, subjects fasted for 5 hours before each of 14 days of training sessions and 3 days of test sessions. Subjects were rewarded with water for accurate trial responses. During the test sessions, BOLD signals were recorded from regions including VS and dACC. Although BOLD signals in both areas were associated with positive and negative RPEs, only those in dACC associated with negative RPE showed a significant correlation with performance improvement. Additionally, no significant correlation was observed between BOLD signals associated with RPEs in VS and dACC. These results suggest that although signals associated with positive and negative RPEs from both midbrain and cortical systems are readily accessible, only RPE signals in the prefrontal system, generated without linking to RPE signals in VS, are utilized for the enhancement of VPL.

## Introduction

Visual perceptual learning (VPL) refers to the process of improving visual performance through repetitive practice or exposure to visual features^1–7^. VPL is primarily associated with alterations in visual processing including area V1 and is essential for understanding visual plasticity^8–11^. Like other types of learning and memory, a key aspect of VPL is its significant improvement through rewards ^12–19^. However, the role of reward prediction error (RPE) in VPL remains unclear.

The concept of reward prediction error (RPE) involves measuring the difference between expected and actual rewards, playing a crucial role in the learning process by means of reinforcement mechanisms^20–23^. Extensive research has demonstrated that neurons in the midbrain dopaminergic systems strongly respond to RPE in conditioning contexts^24–26^.

Initially, RPE was linked to phasic and rudimentary associative learning processes in the midbrain systems^24^. However, recent studies have unveiled sustained RPE-related responses in prefrontal regions in complex environmental frameworks^27^. Notably, the dorsal anterior cingulate cortex (dACC), traditionally known for error detection through accuracy feedback^28^, which has emerged as a contributor to RPE processing^29–33^.

Yet, the specific interaction between RPE signals in the midbrain dopaminergic systems and those in the prefrontal systems, potentially functioning as discrete entities, to enhance learning during training, remains unclear.

To address this question, we conducted orientation detection task training with participants. Each correct response in the trials was rewarded with a drop of water, as illustrated in Fig. 1a. Our focus was on analyzing Blood Oxygenation Level Dependent (BOLD) signals in three specific brain regions: the ventral striatum (VS), dACC, and V1. These signals were measured before training, one day after the initial training, and one day after completing the training. Our aim was to examine the presence of RPE-related signals in these regions and their role in the development of VPL.

**Fig. 1:**
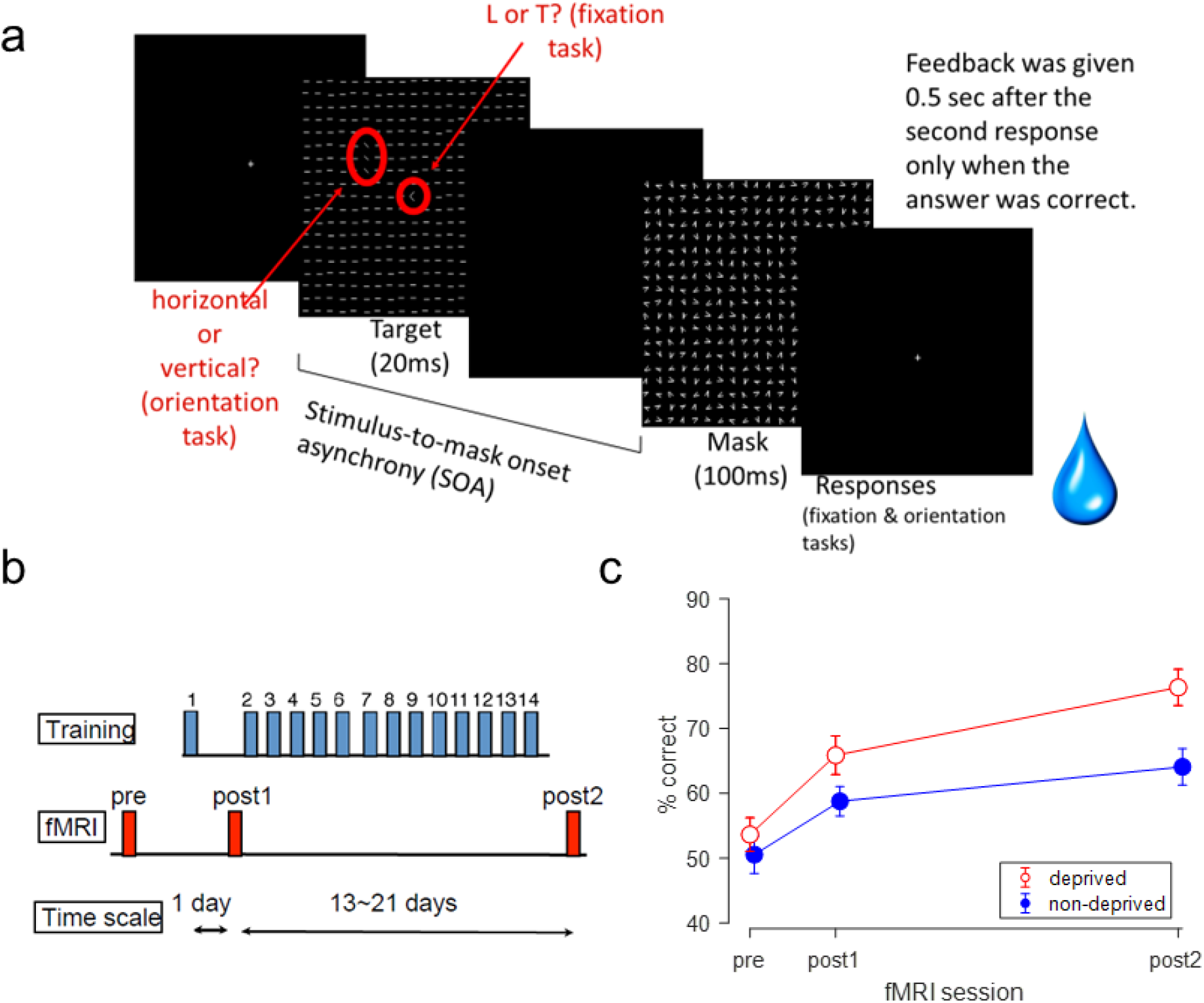
Overview of Texture Discrimination Task, Training Protocol, and Performance Outcomes. a. Each trial’s procedure in the texture discrimination task. b. The full study comprised 14 days of training sessions and 3 days of fMRI and testing sessions. c. Mean performance (± standard error) in the main experiement under deprived condition (red open circles) and the control experiment under non-deprived condition (blue filled circles) across pre, post 1 and post 2 fMRI and test sessions.

**Fig. 2:**
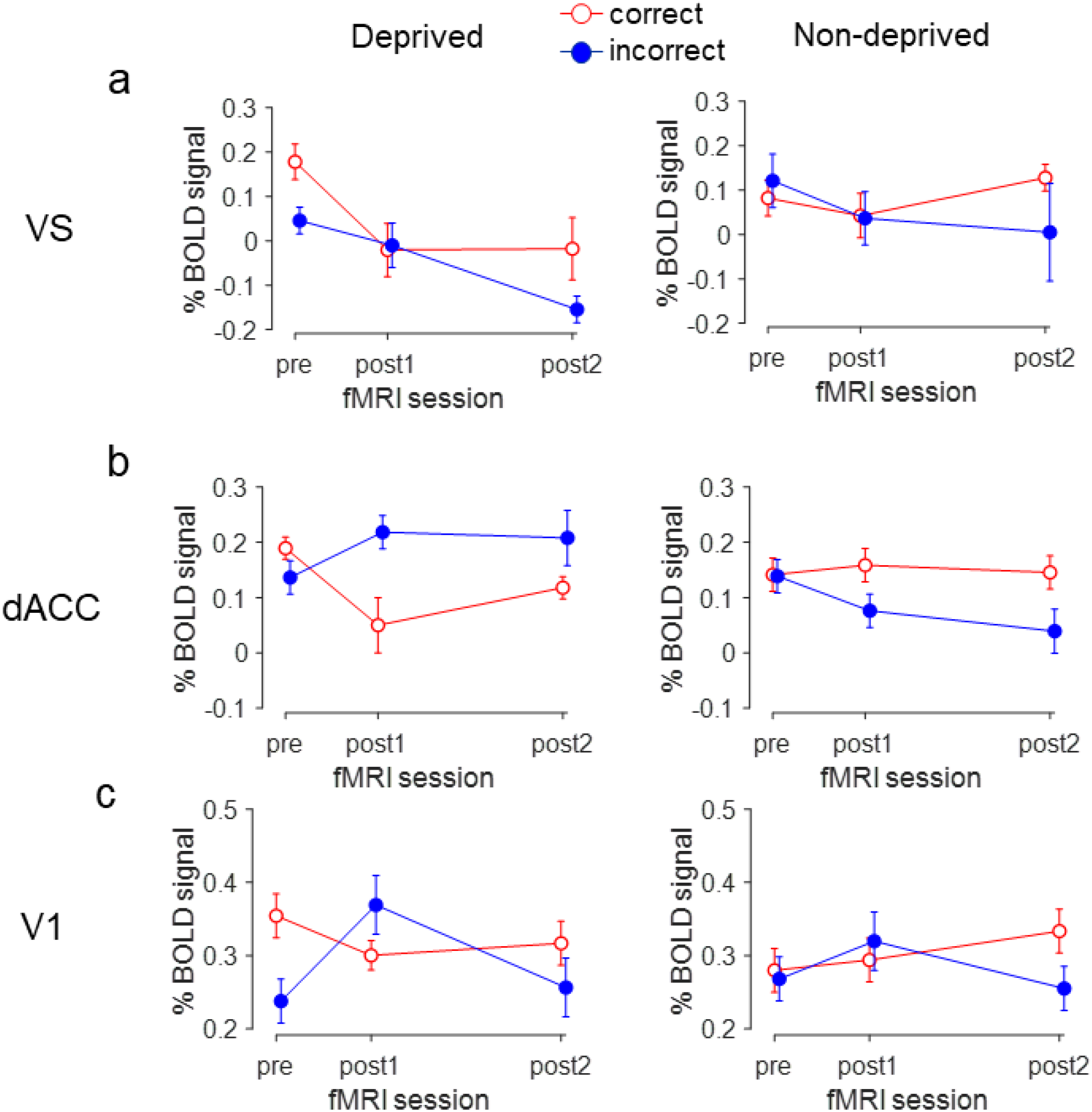
Training-Stage Dependent BOLD Signal Changes in VS, dACC, and V1 under Deprived (Main) and Non-Deprived (Control) Expriments. The mean percentage of BOLD signals (± standard error) as a function of the training stage for deprived (left) and non-deprived (right) experiments in VS (a), dACC (b) and V1 (c). The zero point in % BOLD signal change represents no deviation from the baseline, which is the average BOLD signal measured at the first two time points (TRs).

We observed that signal patterns in both VS and dACC reflect RPE. However, only the signals in dACC exhibited significant correlations with performance improvements. Moreover, there is no significant correlation found between signals for negative RPEs in VS and dACC. These findings suggest that while both midbrain and prefrontal systems generate readily accessible RPE signals during VPL training, the RPE signals in dACC are not directly influenced by RPE signals in VS. Instead, these signals in dACC significantly contribute to enhancing VPL.

## Results

To explore the role of RPE in VPL, we conducted an experiment using psychophysical and brain imaging methods. We tracked BOLD signal changes in VS, dACC and V1 in three stages, aiming to investigate the relationship between activations in these areas and positive and negative RPEs. We also examined whether activation related to RPE contributes to performance improvement and examine potential associations among RPE signals in different areas. The entire experiment comprised 14 days of training and 3 days of fMRI and test sessions. Throughout both training and test sessions, we measured brain activations using fMRI, while the performance in the trained task was evaluated using psychophysics, as illustrated in Fig. 1b. Given that previous studies have shown significant VPL after the first day of training^34,35^, fMRI and test sessions were conducted not only before and after the training but also one day after the first training session^34^.

In each trial of both the training and fMRI and test sessions, 15 subjects (6 females) were asked to perform a texture discrimination task (TDT)^35^, which is a standard method for VPL training (Fig. 1a). For the TDT, subjects were first asked to identify whether the letter at the center of the stimulus was an “L” or a “T” to ensure that they maintained their gaze at the display center. Subsequently, the subjects were asked to report whether the orientation of the array of 3-line segments differing in orientation from the other segments and presented in the upper left quadrant of the stimulus was horizontal or vertical. Subjects were instructed not to consume food or drinks for 5 hours before each daily training or test session. At the beginning of each training or test session, subjects were asked to have a tube connected to a water feeder in their mouths. Upon a correct response to the second task, a drop of water was dispensed into their mouths, while no water was given for incorrect responses. Our prior studies have demonstrated that a drop of water functions effectively as a reward (cite our papers using water as reward).

During the test sessions, trial procedures were identical to those of the training sessions, with one exception: while the training sessions were conducted in a small experimental room, test sessions were conducted with subjects lying inside the scanner to measure BOLD signals in the regions of interest.

### 1. Performance improvements and RPE

We examined whether performance improvement were observed in this experiment. What would the potential performance improvement mean for positive and negative RPEs? Positive RPE should occur in the situation in which reward is given when subjects thought of their response as incorrect. The performance improvement increases the predictability of correct responses and should decrease positive RPE. In contrast, negative RPE should occur in the situation in which reward is not given despite the fact that subjects regarded their response as correct. The performance improvement decreases the predictability of incorrect responses. When subjects receive no reward due to incorrect response, strong negative RPE should occur.

Fig. 1c shows the percentage of correct responses as a function of the test session stage. We conducted a one-way ANOVA with “Session” (pre, post 1, vs. post 2) as the within-subject factor. Our analysis revealed significant effect of Session [F(2,28) = 16.048, p < 0.001, partial η^2^ = 0.534, 95% confidence interval CI = (0.348, 0.696)]. Upon these results, we further conducted post hoc tests utilizing simple effects analysis approach. These results indicate a significant increase in performance.

Since significant performance increase occurred in this experiment, as learning progresses positive RPE should decreases while negative RPE should increase in both conditions.

### 2. BOLD signal related to positive or negative RPE

For the next step, we examined whether any BOLD signal in VS, dACC, and V1 were associated with positive or negative RPE. As mentioned above, our results indicate that performance tended to increase over training. In this case, positive RPE should decrease while negative RPE should increase as learning progresses. BOLD signal changes in brain areas corresponding to correct responses as learning progresses are regarded as related to negative RPE, whereas those for incorrect responses related to positive RPE^36–38^.

#### 2-1. Prediction error in VS

For data analysis, we first conducted a mixed two-way model ANOVA with the within-subject factors of Session (pre, post 1 vs. post 2) and Response (correct vs. incorrect). The results showed a significant main effect of Session [F(2,28) = 6.691, p = 0.004, partial η^2^ = 0.323, CI = (0.144, 0.527)]. On the other hand, we did not find a significant effect of Response [F(1,14) = 3.246, p = 0.093] or interaction between Session and Response [F(2,28) = 2.113, p = 0.140].

These results indicate decreased activations in VS with decreasing positive and negative RPEs as learning progressed. They are in accordance with previous research that VS increases the activity with stronger positive RPE, whereas it decreases the activity with stronger negative RPE^20,24,38,39^.

#### 2-2. Prediction error in dACC

Following a similar methodology applied to BOLD signals in VS, we conducted a two-way ANOVA with Session (Pre, Post 1 vs. Post 2) and Response (correct vs. incorrect). The results revealed a significant interaction between Session and Response [F(2,28) = 5.674, p = 0.009, partial η^2^ = 0.288, CI = (0.117, 0.496)], but no significant main effects for either session [F(2,28) = 0.885, p = 0.424] or response [F(1,14) = 2.532, p = 0.134]. Because of the significant interaction, we conducted a simple effects analysis. For correct responses, BOLD signals significantly decreased from the pre-test to post 1 test [t(14) = 2.904, p = 0.012, Cohen’s d = 0.750, CI = (0.164, 1.316)]. Conversely, for incorrect responses, BOLD signals exhibited a significant increase from pre-test to post 1 test [t(14) = −2.435, p = 0.029, Cohen’s d = −0.629, CI = (−1.176, −0.063)].

These results indicate that for the correct responses, dACC exhibited decreased activation from pre-test to post 1 test, correlating with decreasing positive RPE. A similar pattern was observed in VS. In contrast, for the incorrect responses associated with negative RPE, dACC activation tended to increase with increasing negative RPE, particularly noticeable in the early phase of training, wherein performance increase tended to be more pronounced than in the later stage.

The increased activation in dACC with increasing negative RPE contrasts with the decrease in activation observed in VS. However, this increased activation is in accordance with previous studies that suggest that dACC response reflects surprise, consequently leading to attention-driven learning^33^.

#### 2-3. Prediction error reflected in V1

As VPL is likely to be associated with changes in V1^6,34,40,41^, we also analyzed BOLD signals in V1. The results of a two-way ANOVA with Session (Pre, Post 1 vs. Post 2), Response (correct vs. incorrect) showed a significant interaction between Session and Response [F(2,28) = 6.872, p = 0.004, partial η^2^ = 0.329, CI = (0.149, 0.532)]. However, there were no significant main effects for either Session [F(2,28) = 2.096, p = 0.142] or Response [F(1,14) = 0.891, p = 0.361]. Because of the significant interaction, we further simple effects analysis in a similar way as in dACC. For correct responses, BOLD signals in V1 significantly decreased from the pre-test to the post 1 test [t(14) = 2.804, p = 0.014, Cohen’s d = 0.724, CI = (0.142, 1.286)]. Conversely, for the incorrect responses, BOLD signals in V1 significant increased from the pre-test to post 1 test [t(14) = −2.823, p = 0.014, Cohen’s d = −0.729, CI = (−1.291, −0.146)].

These results indicate that activation in V1 decreased as positive RPE decreased and increased as negative RPE increased, as learning progressed. This pattern of results is basically the same as those in dACC.

### 3. Prediction error signals and performance improvements

The above results suggest that RPE signals are readily accessible in VS, dACC and V1, although there is an opposite trend in the signs (increase vs. decrease) for negative RPE between VS and dACC & V1. This leads us to explore whether RPE signals from all these areas are effectively utilized for learning. To address this, we investigated the correlation between changes in BOLD signals in VS, dACC and V1 and the increasing performance across subjects.

For VS, no significant correlations were observed between performance change and the BOLD signal for either correct responses (post 1 - pre: R = 0.031, p = 0.912; post 2 - pre: R = −0.369, p = 0.177) or incorrect responses (post 1- pre: R = −0.156, p = 0.578; post 2 - pre: R = 0.15, p = 0.594) (Fig. 3a). These results suggest that BOLD signal in VS associated with positive and negative RPE shown above do not contribute to VPL.

**Fig. 3:**
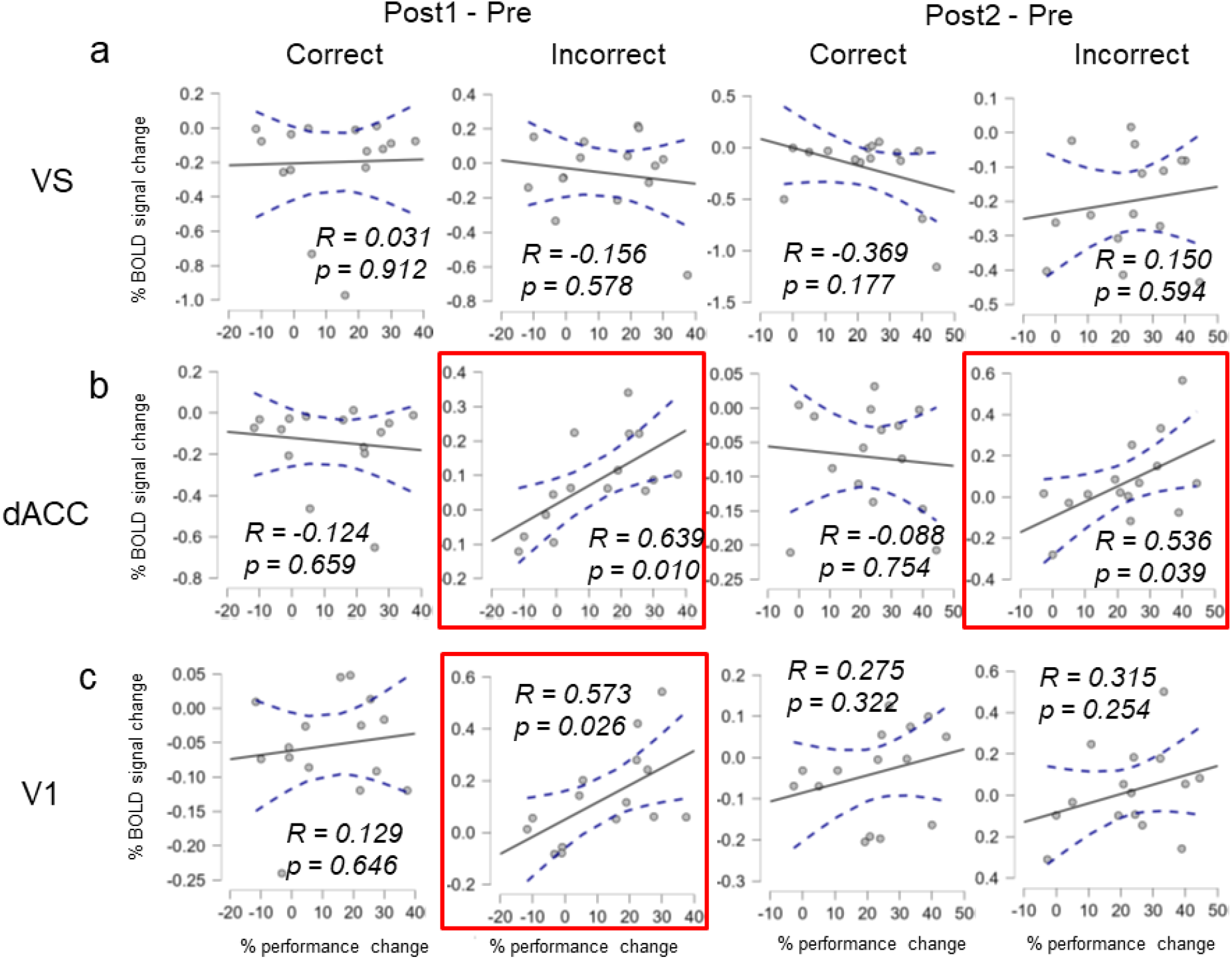
Correlation of Performance Improvement with BOLD Signal Changes in VS, dACC, and V1 during Deprived (Main) Experiment. The correlation between performance and activation changes in VS (a), dACC (b), and V1 (c) in the deprived condition. This analysis is focused on correct and incorrect trials in the deprived condition. On the x-axis, we have performance change, while the y-axis represents the percentage change in BOLD signal observed between sessions. Dashed lines represent the 95% confidence interval. The first two columns present correlations during the early learning phase (post 1 - pre), while the subsequent two columns showcase correlations across the entire learning trajectory (post 2 - pre).

**Fig. 4:**
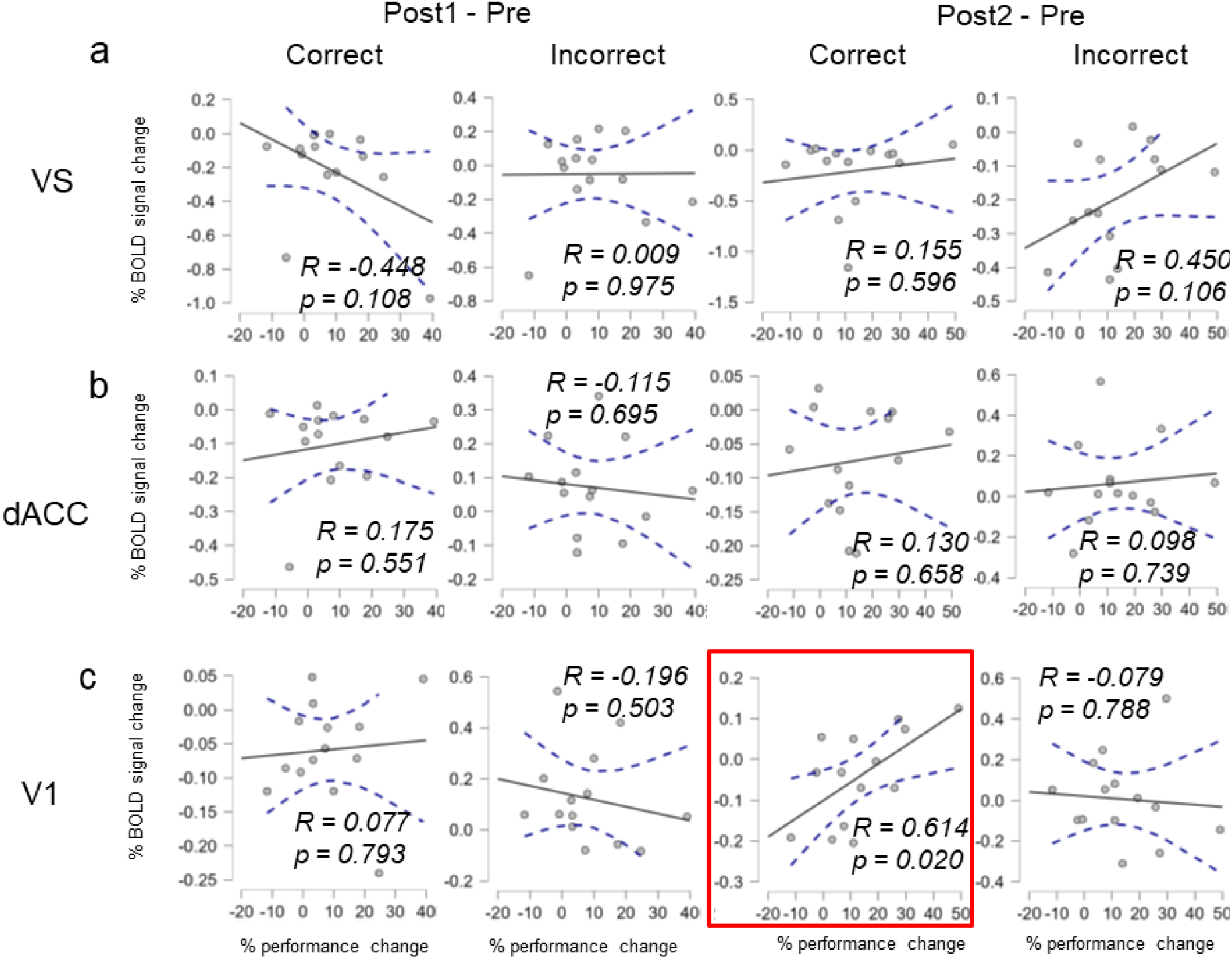
Correlation of Performance Improvement with BOLD Signal Changes in VS, dACC, and V1 during Non-Deprived (Control) Experiment. The correlation between performance and activation in VS (a), dACC (b), and V1 (c) in the deprived condition. This analysis is focused on correct and incorrect trials in the deprived condition. The x-axis represents the percentage of performance change, while the y-axis represents the percentage in BOLD signal change observed between sessions. Dashed lines represent the 95% confidence interval. The first two columns present correlations during the early learning phase (post 1 SNR - pre), while the subsequent two columns showcase correlations across the entire learning trajectory (post 2 - pre).

In contrast, for dACC, we identified a significant positive correlation between performance changes and BOLD signal changes for incorrect responses in both the early (post 1 - pre: R = 0.639, p = 0.010) and across the overall learning progress (post 2 - pre: R = 0.536, p = 0.039) (Fig. 3b). However, we observed no significant correlation between performance changes and BOLD signal changes for correct responses either in the early (post 1 - pre: R = −0.124, p = 0.659) or across the overall learning progress (post 2 - pre: R = −0.088, p = 0.754). These results suggest that signals in dACC associated with negative RPE play a significant role in VPL.

With regard to V1, we found a significant positive correlation between performance changes and BOLD signal changes for incorrect responses that was associated with negative RPE during the early learning phases (post 1 - pre: R = 0.573, p = 0.026) (Fig. 3c). However, no significant correlation was observed between performance changes and BOLD signal changes for correct responses either in the early (post 1 - pre: R = 0.129, p = 0.646) or across the entire learning progression (post 2 - pre: R = 0.275, p = 0.322).

### 4. Relationship between negative prediction error signals in VS and dACC

Given the observed significant BOLD signal changes —albeit with opposite signs— associated with negative RPE in both VS and dACC, it is crucial to investigate their potential relationship. The correlation analysis from the above tests suggests that negative RPE signal in VS does not significantly contribute to VPL. Nevertheless, this result does not rule out the possibility that negative RPE signal in VS might indirectly impact VPL by directly influencing the generation of negative RPE signal in dACC. If this were the case, one would expect a negative or positive correlation between BOLD signal changes in VS and dACC. However, we found no significant correlation between BOLD signal changes in these areas either from pre- and post 1 stages (R = 0.323, p = 0.240) or from pre-and post 2 stages (R = 0.261, p = 0.347). This result does not support the hypothesis that activation related to the negative RPE signal in VS directly influences the generation of the negative RPE signal in dACC.

#### Control experiment

Water in the above-mentioned main experiment might have functioned as a reward. However, it could have also served as accuracy feedback, which is known to enhance learning^42,43^. To determine which is the case, we conducted a control experiment to assess the value of water as a reward. In this experiment, a new group of subjects (n= 14, 5 females) was allowed to eat or drink as usual. If the amplitude of VPL in this control experiment is similar to that in the main experiment, it suggests that accuracy feedback was the primary driver of VPL. Conversely, if the amplitude of VPL in the control experiment is significantly lower than that in the main experiment, it suggests that water, functioning as a reward, substantially contributes to VPL.

Fig. 1c shows performance improvements in the main and control experiments. We conducted two-way ANOVA on performance with “Session” (pre, post 1, vs. post 2) as the within-subject factor and “Experiment” (main vs. control) as the between-subject factor. We obtained significant effects of both Session [F(2,54) = 21.902, p < 0.001, partial η^2^ = 0.448, CI = (0.308, 0.582)] and Condition [F(1,27) = 5.054, p = 0.033, partial η^2^ = 0.158, CI = (0.004, 0.429)]. However, there was no significant interaction between Session and Condition [Session x Condition: F(2,54) = 1.397, p = 0.256]. These results suggest that both the water presence in the deprived situation of the main experiment and the non-deprived situation in the control experiment drove VPL. However, the VPL magnitude in the deprived situation was significantly larger than in the non-deprived situation. Consequently, we performed one-way ANOVA selection using Session as the within-subject factor on the results of the control experiment. The results revealed a significant main effect of Session [F(2,28) = 6.734, p = 0.004, partial η^2^ = 0.325, CI = (0.146, 0.529)]. These results are in accord with the hypothesis that water significantly drives VPL. Additionally, these results suggest that, whether or not subjects were deprived of water or food, positive RPE decreased while negative RPE increased with more training, although the magnitudes of the changes in RPEs should be smaller without deprivation.

We then tested whether BOLD signals in VS, dACC and V1 are related to positive and negative RPE in the control experiment in which no fasting occurred with subjects. We conducted a two-way ANOVA with the main factors being Session (pre, post 1 vs. post 2) and Response (correct vs. incorrect) to the BOLD signals in VS, dACC and V1 measured in the control experiment. In VS, we observed no significance in any of the main effect of Session [F(2,26) = 0.499, p = 0.613], the main effect of Response [F(1,13) = 0.500, p = 0.492] and interaction between Session and Response [F(2,26) = 0.799, p = 0.461]. In dACC, there was a significant main effect of Response [F(1,13) = 4.874, p = 0.046, partial η^2^ = 0.273, CI = (0.008, 0.639)]. However, there was no significant main effect of Session [F(2,26) = 1.717, p = 0.199] or interaction between Session and Response [F(2,26) = 1.079, p = 0.355]. In V1, there was no significant main effect of Session [F(2,26) = 0.924, p = 0.410] or Response [F(1,13) = 0.306, p = 0.590] or interaction between Session and Response [F(2,26) = 1.271, p = 0.298].

These results suggest that no signals in any of the three areas are significantly associated with either positive or negative RPE. Given that water in the control experiment was effective either only for accuracy feedback or for both accuracy feedback and a smaller reward value than in the main experiment, accuracy feedback alone is insufficient to generate signals associated with RPE in any of these areas. In other words, a reward is necessary for generating signals associated with RPE in these areas.

## Discussion

This study aims to explore the utilization of RPE signals from the VS in midbrain systems and the dACC in prefrontal systems in reinforcing and enhancing VPL. First, we found that RPE signals were generated in these distinct systems. During training, we found that as learning progressed, BOLD signals in the VS decreased for both correct and incorrect responses. These results are in accordance with previous research in associative learning, indicating that VS responses increase with positive RPE but decrease with negative RPE^24,44,45^. On the other hand, as learning progressed, BOLD signals in the dACC increased for incorrect responses with no significant changes observed for correct responses. The results for incorrect responses are in accordance with earlier studies that negative RPE is associated with increased responses in dACC^33^. Second, the observed pattern of BOLD signals in V1 was consistent with that in dACC, suggesting significant involvement of cortical areas in reward-related enhancement of VSS. Third, no significant correlation was found between the changes in BOLD signals in VS associated with positive and negative RPEs and the performance improvement. In contrast, the <u>increased</u> BOLD signals in dACC associated with negative RPE were significantly correlated with enhanced performance improvement. Moreover, no significant correlation was observed between the BOLD signal changes associated with negative RPE in VS and those in dACC. These results suggest that negative RPE signal in VS does not directly influence that in dACC or VSS.

The present results align, at least partially, with recent research indicating that both VS and dACC play roles in PREs, although their functions in learning are distinct. Namely, VS activity depends on the sign of PRE. It is enhanced with positive PRE, while being suppressed with negative PRE. This pattern of activity aligns with the RPE principles of reinforcement learning^24,44,45^. On the other hand, the dACC exhibits <u>increased</u> activity in response to negative PRE^33^. This enhanced activity is consistent with the error-related negativity (ERN), a component of the event-related brain potential that emerges in the region including dACC when human participants make errors in tasks (cite the papers about ERN). The increased activity associated with negative RPE may be driven by attention due to the surprise elicited by error ^46–51^. Indeed, these two distinct roles in the midbrain including the VS and frontal systems including the dACC are widely accepted^22,52–56^.

At the same time, the absence of a significant correlation between BOLD signal changes in VS for positive or negative RPEs and the increase in performance suggests that signals associated with neither positive nor negative prediction error in VS are utilized for VPL, in spite of the fact that these signals are ready to be used. In contrast, the significant positive correlation between BOLD changes in dACC associated with negative RPE and increase in performance suggests that increased signals associated with negative RPE in dACC are indeed utilized for VPL.

How do signals associated with negative prediction error in the midbrain system relate to those in the prefrontal system? The lack of a significant correlation between BOLD signal changes associated with negative RPE in the VS and dACC suggests that VPL is facilitated by increased activity associated with negative RPE in the dACC without direct reliance on RPE signals in VS. However, it cannot be ruled out that signals in the midbrain, apart from those associated with RPE, might contribute to the enhancement of signals associated with negative RPE in VS ^32,55^. Future research needs to explore the validity of this possibility.

It is noteworthy that our results showed no evidence for activation in any of the three areas associated with either positive or negative in the non-deprived condition. As accuracy feedback was provided in the control experiment, the absence of significant activation associated with negative RPE suggests that accuracy feedback, in the absence of reward, is insufficient to generate negative RPE signals.

What is the role of V1? Studies have shown significant involvement of V1 in VPL^6,34,40,41^. V1 is more likely to undergo modifications that enhance sensitivity to the trained feature rather than being directly involved in generating signals for reinforcement or attention. Our results, indicating a correlation between V1 signals associated with negative RPE with performance improvement, may suggest that these signals emanate from areas including the dACC and are transmitted to V1.

Why do dACC signals impact VPL while VS signals do not? The VPL observed in this study’s experiments falls under the category of task-relevant VPL, in which VPL of a visual feature of a trained task occurs^1,57^. Task-relevant VPL involves various cortical areas, including cognitive areas^13,34,41,58–60^, and visual areas including the V1 which are changed to retain VPL for a long time^34^. Although the VS is typically linked to simple associative learning predicted on reward contingencies^57,61^, our findings suggest that the task at hand engages more complex cognitive processes, as demonstrated by the activation of cortical areas such as the dACC and V1 in response to negative RPE. We observed that as learning progressed, BOLD signals in both the dACC and V1—cortical areas—increased with negative RPE, contrasting with the VS where the BOLD signal decreased. The increased activation in the dACC and V1 can be attributed to negative RPE eliciting attention^33,48–51^, which in turn activates attentional source areas in prefrontal regions, including the dACC. This attention may then lead to both activation and long-term network modifications within areas involving V1^34^.

This study investigated how activity in the VS and dACC associated with positive and/or negative RPE, occurs and influences VPL. In our experiment, we observed that activation in the VS increased with positive RPE and decreased with negative RPE. Conversely, in the dACC, we found that signals related to negative RPE were not decreased but increased. However, only these increased signals in dACC were significantly correlated with performance improvement. Moreover, no significant correlation was observed in activations between the VS and those in cortical areas such as the dACC and V1. These results suggest that although RPE signals in both the VS and dACC are computed and ready to be used, only the signals in dACC, which are generated without direct interactions with RPE signals in VS, are utilized to enhance VPL.

## Methods

### Subjects

In this study, 29 healthy adults (11 females and 18 males) aged 18 to 39 years participated, with a mean age of 25.62 years ± 5.79 standard deviation (SD). The participants had normal vision or corrected-to-normal vision, and 15 of them (6 females, mean age 25.26 years ± 5.58 SD) fasted for five hours before the experiment (high reward value condition), while the other 14 participants (5 females, mean age 26 years ± 6.2 SD) did not fast (low reward value condition). All subjects gave written informed consent for the experimental protocol, which was approved by the Institutional Review Board.

### Behavioral Training Session

The experiment was held in a dimly lit room with a CRT monitor with a 19-inch screen,1024×768 pixel resolution, and 100 Hz refresh rate. The distance between the screen and the participant’s eyes was 57 cm. Throughout the experiment, the tube was held in the mouth to receive water. The study utilized the widely used Texture Discrimination Task (TDT)^35^. During each trial, a task screen would appear for 17 ms, followed by a blank screen with varying duration, then a mask screen with randomly placed V-shaped patterns for 100 ms. The shorter the time between the task screen and mask screen onset (known as the stimulus-to-mask-onset asynchrony or SOA), the harder the task. The task screen consisted of horizontal bars with a letter (’T’ or ‘L’) at the center and a target array of three diagonal bars arranged in a horizontal row or vertical column, located in the upper left or right visual field. The location of the target array was randomly selected and provided to each participant. The participants were asked to perform two tasks during the experiment, starting with identifying the letter at the center of the screen in the fixation task and then determining the orientation of the target array in the orientation discrimination task, using two buttons on a keypad. They received immediate auditory feedback for their fixation task response, with a high-pitched sound indicating a correct answer and a low-pitched sound signaling an incorrect response. Feedback for the orientation task in the form of water was only given if the response was correct and it was provided 500 ms after the second response^12,41^. Water was delivered for 200 ms through a plastic tube using a ValveLink®8.2system made by Automate Scientific, Inc^12^. Each training session consisted of 6 blocks with 120 trials in each block, for a total of 720 trials. The SOA became shorter as training progressed, and in the first session, SOAs of 500ms, 250ms, 200ms, 150ms, 100ms, and 67ms were applied to all participants in sequence, from the first block to the last. In subsequent sessions, an optimized SOA list was generated for each individual, based on their performance in the previous session, using curve fitting with a logistic function.

### fMRI Experiments

The MRI scan was conducted in three separate sessions that were not scheduled in conjunction with the training sessions. The first session (pre) occurred at least one day prior to the initial training session, the second session (post1) happened precisely one day following the first training session, and the third session (post2) took place at least one day after the last training session. Participants completed the task while lying in the MRI scanner and looking at a mirror that projected stimuli through an LCD projector (1024 x 768, 75 Hz). Throughout the experiment, they held a tube in their mouth to drink water. Although the task was the same as the training session, a single SOA (100ms) was used for comparing the different sessions. The timing between each trial was randomized using optseq2 software to enhance statistical efficacy in the event-related fMRI paradigm. A total of 120 trials were conducted, distributed over four runs (30 trials per run), to minimize the learning effect in the MRI session. Each trial lasted up to four seconds, and one run consisted of a task period of 120 seconds and a rest period of 120 seconds, totaling 240 seconds.

### Image Acquisition

The Participants in the study were scanned using a 3T MR scanner, either a Trio or Prisma (both from Siemens), with a head coil in place for the entire duration of the experiment. During the study, the MR scanner was upgraded from a Trio to a Prisma, but the same protocol was followed as closely as possible. BOLD contrast was measured using a gradient-echo EPI sequence with TR=2s, TE=28ms, and flip angle=90 degrees. Thirty-three slices were acquired, each 3mm x 3mm x 3mm and parallel to the AC-PC plane, to cover the entire brain. For anatomical reconstruction, three T1-weighted MR images were obtained using MPRAGE with TR=2.3s, TE=2.28ms, flip angle=8 degrees, TI=900ms, 256 slices, and a voxel size of 1mm x 1mm x 1mm.

### Image Pre-processing

We began by concatenating the EPI runs before preprocessing. The fMRI data was processed with FEAT, which is part of the FSL library. To register T1-weighted images to high-resolution structural and/or standard space images, we used FLIRT^62,63^, and further refined the registration from high resolution structural to standard space using FNIRT nonlinear registration^64,65^. For preprocessing the EPI images, we utilized FSL, including correction for slice timing and motion using MCFLIRT^66^, spatial normalization to the MNI standard space using the resampled T1 image with 2mm voxel size, and spatial smoothing using a 5mm Gaussian kernel.

### fMRI data analysis

In this study, the fMRI data were analyzed using Region of Interest (ROI) analysis to investigate specific brain regions^67–69^. The ROIs for V1, dACC, and the striatum were created using FreeSurfer’s anatomy masks^70^. The V1 ROI was defined by the trained hemisphere, and the dACC and ventral striatum ROIs used both hemispheres. The dACC mask was formed by combining parcels a32pr and p32pr, using an individualized HCP-MMP parcellation generated with FreeSurfer^71^. The VS ROI was created by combining anatomical masks and functional activation maps. GLM analysis was performed on the pre-test fMRI data from the deprived and non-deprived groups^72,73^. The analysis modeled two active conditions against the baseline, with the onset times of correct and incorrect trials. The beta maps resulting from the analysis identified significant clusters that underwent second-level group analysis. The group beta map was transformed to the individual T1 space, and an individualized ventral striatum ROI was generated by overlapping the NAc, putamen, and caudate regions from FreeSurfer with the significant beta map. To calculate the percentage change in BOLD signal within these ROIs, the BOLD signal intensity was measured at each time point in the fMRI time series. The percentage signal change was calculated by subtracting the baseline signal intensity (average of the first and second TRs) from each time point, dividing by the baseline intensity, and then multiplying by 100^67–69^. For correct trials, the mean BOLD signal for the third TR (associated with visual stimulation, accounting for a 6-second BOLD delay) and the fourth TR (associated with water feeding) were averaged. For incorrect trials, only the mean BOLD signal for the fourth TR was taken into account, as no water was provided. We specifically focused on prediction errors for incorrect trials, where participants expected a reward but did not receive one. Therefore, it is essential to analyze the BOLD signal corresponding to the 4th TR, which coincides with the time when the water feeding was the anticipated but do not occur. The 3rd TR was excluded from this analysis for incorrect trials because it was associated with the stimulus presentation and occurred before the ‘regret’ point—the moment participants realized their response was incorrect—which occurred at least 0.5 seconds after the second response, more than 1.5 seconds after the stimulus onset (3rd TR). For these reasons, the BOLD signal from the 3rd TR was not considered for incorrect trials. Open red circles indicate correct trials, while filled blue circles indicate incorrect trials. Error bars represent the standard error.

### Statistical analysis

A mixed model ANOVA, supplemented by post hoc assessments through simple effects analysis, was conducted using JASP 0.17.3^74^. To caculuate standard error for repeated measure ANOVA, we applied the method by Morey (2008)^75^, which is implemented in JASP.

---

**Supplementary Fig. 1:**
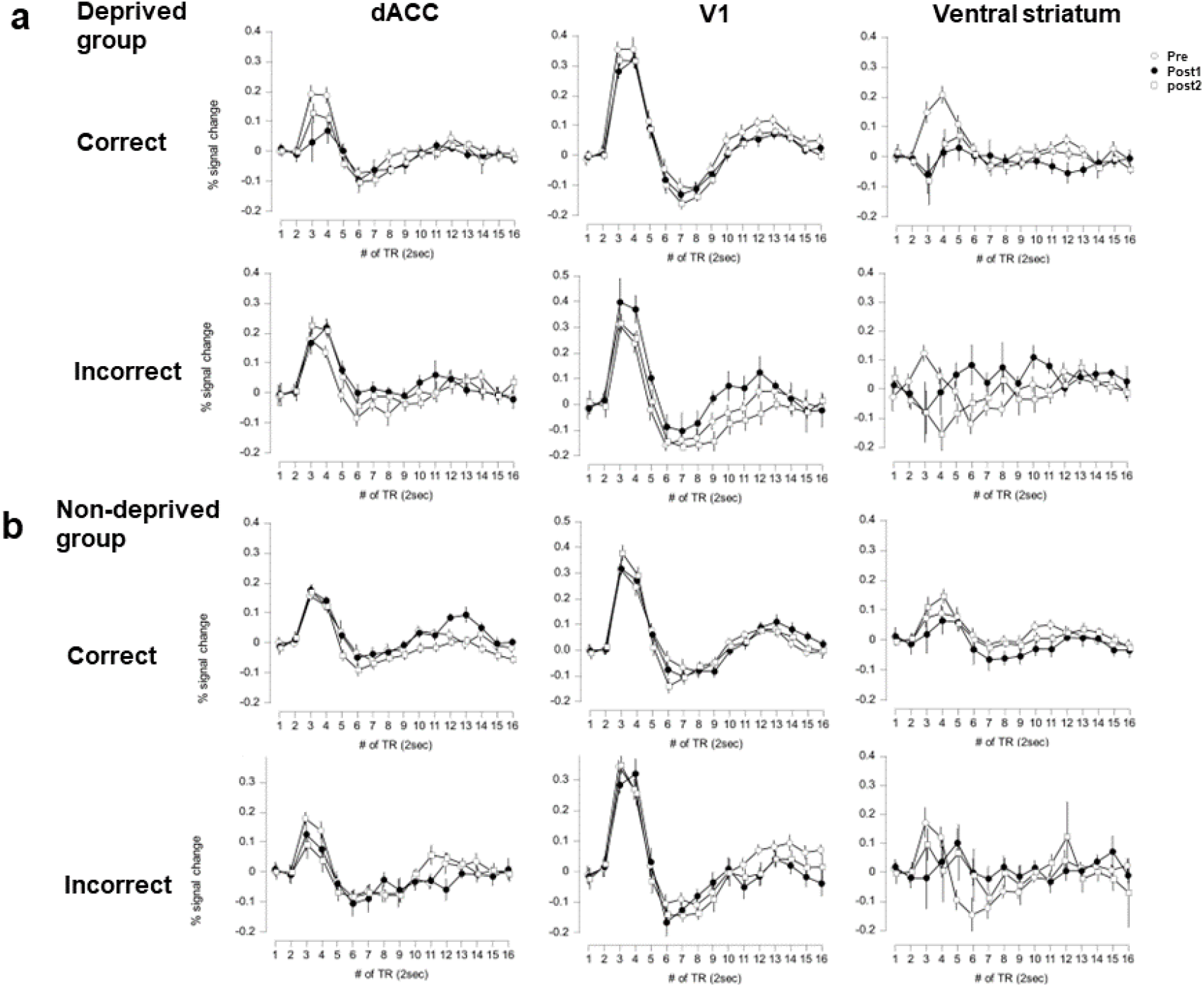
BOLD Signal Changes in dACC, V1, and VS Across Pre-, Post-1, and Post-2 Tests. This figure shows BOLD signal changes in three brain regions: the dACC, V1, and VS. The x-axis shows time in time repetitions (TRs), each lasting 2 seconds. BOLD signal intensity was measured at each time point, and the percentage change from baseline (average of the first and second TRs) was calculated by subtracting this baseline, dividing by it, and multiplying by 100. Open circles represent the mean percentage change in the pre-test, filled circles for the post-1 test, and open squares for the post-2 test, with error bars indicating standard errors. Panel a depicts the main experiment with the deprived group, and panel b shows the control experiment with the non-deprived group, with correct and incorrect trials presented separately for each.

## Acknowledgements

This research was supported by grants from NIH (R01 EY031705, R01 EY027841, R01 EY019466), National Research Foundation of Korea (2016R1C1B2015901), and Institute for Basic Science, Korea (IBS-R015-D1). The authors would also like to extend their gratitude to HyungGoo Kim and Min-Suk Kang for their comments.

## Author contributions

DK, YS and TW designed the experiment. DK conducted the experiments and analyzed the data. DK and TW wrote the original draft. All authors were involved in reviewing and editing.

## References

1 Sasaki, Y., Nanez, J. E. & Watanabe, T. Advances in visual perceptual learning and plasticity. Nat Rev Neurosci 11, 53–60 (2010). 10.1038/nrn2737

2 Lu, Z.-L. & Dosher, B. A. Current directions in visual perceptual learning. Nature Reviews Psychology 1, 654–668 (2022).

3 Watanabe, T., Nanez, J. E. & Sasaki, Y. Perceptual learning without perception. Nature 413, 844–848 (2001). 10.1038/35101601

4 Seitz, A. R. & Watanabe, T. Psychophysics: Is subliminal learning really passive? Nature 422, 36 (2003).

5 Shibata, K., Watanabe, T., Sasaki, Y. & Kawato, M. Perceptual Learning Incepted by Decoded fMRI Neurofeedback Without Stimulus Presentation. Science 334, 1413–1415 (2011). 10.1126/science.1212003

6 Shibata, K. et al. Overlearning hyperstabilizes a skill by rapidly making neurochemical processing inhibitory-dominant. Nat Neurosci 20, 470–475 (2017). 10.1038/nn.4490

7 Tamaki, M. et al. Complementary contributions of non-REM and REM sleep to visual learning. Nat Neurosci 23, 1150–1156 (2020). 10.1038/s41593−020−0666-y

8 Yu, Q., Zhang, P., Qiu, J. & Fang, F. Perceptual Learning of Contrast Detection in the Human Lateral Geniculate Nucleus. Curr Biol 26, 3176–3182 (2016). 10.1016/j.cub.2016.09.034

9 Dosher, B. A. & Lu, Z. L. Perceptual learning in clear displays optimizes perceptual expertise: learning the limiting process. Proc Natl Acad Sci U S A 102, 5286–5290 (2005). 10.1073/pnas.0500492102

10 Li, W. Perceptual Learning: Use-Dependent Cortical Plasticity. Annu Rev Vis Sci 2, 109–130 (2016). 10.1146/annurev-vision-111815-114351

11 Maniglia, M. & Seitz, A. R. Towards a whole brain model of Perceptual Learning. Curr Opin Behav Sci 20, 47–55 (2018). 10.1016/j.cobeha.2017.10.004

12 Seitz, A. R., Kim, D. & Watanabe, T. Rewards evoke learning of unconsciously processed visual stimuli in adult humans. Neuron 61, 700–707 (2009).

13 Law, C. T. & Gold, J. I. Reinforcement learning can account for associative and perceptual learning on a visual-decision task. Nat Neurosci 12, 655–663 (2009). 10.1038/nn.2304

14 Gold, J. I. & Watanabe, T. Perceptual learning. Curr Biol 20, R46–48 (2010). 10.1016/j.cub.2009.10.066

15 Roelfsema, P. R., van Ooyen, A. & Watanabe, T. Perceptual learning rules based on reinforcers and attention. Trends Cogn Sci 14, 64–71 (2010). 10.1016/j.tics.2009.11.005

16 Kim, D., Ling, S. & Watanabe, T. Dual mechanisms governing reward-driven perceptual learning. F1000Res 4, 764 (2015). 10.12688/f1000research.6853.1

17 Kim, D., Seitz, A. R. & Watanabe, T. Visual perceptual learning by operant conditioning training follows rules of contingency. Vis cogn 23, 147–160 (2015). 10.1080/13506285.2015.1015663

18 Kahnt, T., Grueschow, M., Speck, O. & Haynes, J. D. Perceptual learning and decision-making in human medial frontal cortex. Neuron 70, 549–559 (2011). 10.1016/j.neuron.2011.02.054

19 Arsenault, J. T. & Vanduffel, W. Ventral midbrain stimulation induces perceptual learning and cortical plasticity in primates. Nat Commun 10, 3591 (2019). 10.1038/s41467−019-11527-9

20 Schultz, W. Dopamine reward prediction error coding. Dialogues Clin Neurosci 18, 23–32 (2016).

21 Schultz, W. Predictive reward signal of dopamine neurons. J Neurophysiol 80, 1–27 (1998).

22 Schultz, W. Behavioral theories and the neurophysiology of reward. Annu Rev Psychol 57, 87–115 (2006). 10.1146/annurev.psych.56.091103.070229

23 Bayer, H. M. & Glimcher, P. W. Midbrain dopamine neurons encode a quantitative reward prediction error signal. Neuron 47, 129–141 (2005). 10.1016/j.neuron.2005.05.020

24 Schultz, W., Dayan, P. & Montague, P. R. A neural substrate of prediction and reward. Science 275, 1593–1599 (1997).

25 Nakahara, H., Itoh, H., Kawagoe, R., Takikawa, Y. & Hikosaka, O. Dopamine neurons can represent context-dependent prediction error. Neuron 41, 269–280 (2004). 10.1016/s0896-6273(03)00869-9

26 Bromberg-Martin, E. S., Matsumoto, M. & Hikosaka, O. Dopamine in motivational control: rewarding, aversive, and alerting. Neuron 68, 815–834 (2010). 10.1016/j.neuron.2010.11.022

27 Badre, D. On Task : How Our Brain Gets Things Done. First paperback edition. edn, (Princeton University Press, 2022).

28 Ito, S., Stuphorn, V., Brown, J. W. & Schall, J. D. Performance monitoring by the anterior cingulate cortex during saccade countermanding. Science 302, 120–122 (2003). 10.1126/science.1087847

29 Rushworth, M. F. & Behrens, T. E. Choice, uncertainty and value in prefrontal and cingulate cortex. Nat Neurosci 11, 389–397 (2008). 10.1038/nn2066

30 Matsumoto, M., Matsumoto, K., Abe, H. & Tanaka, K. Medial prefrontal cell activity signaling prediction errors of action values. Nat Neurosci 10, 647–656 (2007). 10.1038/nn1890

31 Brown, J. W. & Braver, T. S. Learned predictions of error likelihood in the anterior cingulate cortex. Science 307, 1118–1121 (2005). 10.1126/science.1105783

32 Holroyd, C. B. & Coles, M. G. H. The neural basis of human error processing: reinforcement learning, dopamine, and the error-related negativity. Psychol Rev 109, 679–709 (2002). 10.1037/0033-295X.109.4.679

33 Hayden, B. Y., Heilbronner, S. R., Pearson, J. M. & Platt, M. L. Surprise signals in anterior cingulate cortex: neuronal encoding of unsigned reward prediction errors driving adjustment in behavior. J Neurosci 31, 4178–4187 (2011). 10.1523/JNEUROSCI.4652-10.2011

34 Yotsumoto, Y., Watanabe, T. & Sasaki, Y. Different dynamics of performance and brain activation in the time course of perceptual learning. Neuron 57, 827–833 (2008). 10.1016/j.neuron.2008.02.034

35 Karni, A. & Sagi, D. The time course of learning a visual skill. Nature 365, 250–252 (1993). 10.1038/365250a0

36 O’Doherty, J. P., Dayan, P., Friston, K., Critchley, H. & Dolan, R. J. Temporal difference models and reward-related learning in the human brain. Neuron 38, 329–337 (2003). S0896627303001697 [pii]

37 Glascher, J., Daw, N., Dayan, P. & O’Doherty, J. P. States versus rewards: dissociable neural prediction error signals underlying model-based and model-free reinforcement learning. Neuron 66, 585–595 (2010). 10.1016/j.neuron.2010.04.016

38 Pagnoni, G., Zink, C. F., Montague, P. R. & Berns, G. S. Activity in human ventral striatum locked to errors of reward prediction. Nat Neurosci 5, 97–98 (2002). 10.1038/nn802

39 O’Doherty, J. et al. Dissociable roles of ventral and dorsal striatum in instrumental conditioning. Science 304, 452–454 (2004). 10.1126/science.1094285

40 Schoups, A., Vogels, R., Qian, N. & Orban, G. Practising orientation identification improves orientation coding in V1 neurons. Nature 412, 549–553 (2001). 10.1038/35087601

41 Yotsumoto, Y. et al. Location-specific cortical activation changes during sleep after training for perceptual learning. Curr Biol 19, 1278–1282 (2009). 10.1016/j.cub.2009.06.011

42 Herzog, M. H. & Fahle, M. The role of feedback in learning a vernier discrimination task. Vision Res 37, 2133–2141 (1997). 10.1016/s0042-6989(97)00043-6

43 Frank, S. M. et al. Efficient learning in children with rapid GABA boosting during and after training. Curr Biol 32, 5022–5030 e5027 (2022). 10.1016/j.cub.2022.10.021

44 Rescorla, R. A. & Wagner, A. W. A theory of Pavlovian conditioning: Variations in the effectiveness of reinforcement and nonreinforcemen. 64–99 (Appleton-Century-Crofts, 1972).

45 Sutton, R. S. & Barto, A. G. Reinforcement learning : an introduction. (MIT Press, 1998).

46 Swan, J. A. & Pearce, J. M. The orienting response as an index of stimulus associability in rats. Journal of Experimental Psychology: Animal Behavior Processes 14, 292 (1988).

47 Courville, A. C., Daw, N. D. & Touretzky, D. S. Bayesian theories of conditioning in a changing world. Trends Cogn Sci 10, 294–300 (2006). 10.1016/j.tics.2006.05.004

48 Holland, P. C. & Gallagher, M. Amygdala circuitry in attentional and representational processes. Trends Cogn Sci 3, 65–73 (1999). 10.1016/s1364-6613(98)01271-6

49 Lee, H. J., Youn, J. M., O, M. J., Gallagher, M. & Holland, P. C. Role of substantia nigra-amygdala connections in surprise-induced enhancement of attention. J Neurosci 26, 6077–6081 (2006). 10.1523/JNEUROSCI.1316−06.2006

50 Belova, M. A., Paton, J. J., Morrison, S. E. & Salzman, C. D. Expectation modulates neural responses to pleasant and aversive stimuli in primate amygdala. Neuron 55, 970–984 (2007). 10.1016/j.neuron.2007.08.004

51 Lin, S. C. & Nicolelis, M. A. Neuronal ensemble bursting in the basal forebrain encodes salience irrespective of valence. Neuron 59, 138–149 (2008). 10.1016/j.neuron.2008.04.031

52 O’Doherty, J. P. Reward representations and reward-related learning in the human brain: insights from neuroimaging. Curr Opin Neurobiol 14, 769–776 (2004). 10.1016/j.conb.2004.10.016

53 Haber, S. N. & Knutson, B. The reward circuit: linking primate anatomy and human imaging. Neuropsychopharmacology 35, 4–26 (2010). 10.1038/npp.2009.129

54 Rushworth, M. F., Kolling, N., Sallet, J. & Mars, R. B. Valuation and decision-making in frontal cortex: one or many serial or parallel systems? Curr Opin Neurobiol 22, 946–955 (2012). 10.1016/j.conb.2012.04.011

55 Cockburn, J. & Frank, M. in Neural Basis of Motivational and Cognitive Control (eds Rogier B. Mars, Jerome Sallet, Matthew F. S. Rushworth, & Nick Yeung) 0 (The MIT Press, 2011).

56 Shenhav, A., Botvinick, M. M. & Cohen, J. D. The expected value of control: an integrative theory of anterior cingulate cortex function. Neuron 79, 217–240 (2013). 10.1016/j.neuron.2013.07.007

57 Watanabe, T. & Sasaki, Y. Perceptual learning: toward a comprehensive theory. Annu Rev Psychol 66, 197–221 (2015). 10.1146/annurev-psych−010814−015214

58 Lewis, C. M., Baldassarre, A., Committeri, G., Romani, G. L. & Corbetta, M. Learning sculpts the spontaneous activity of the resting human brain. Proc Natl Acad Sci U S A 106, 17558–17563 (2009). 10.1073/pnas.0902455106

59 Law, C. T. & Gold, J. I. Neural correlates of perceptual learning in a sensory-motor, but not a sensory, cortical area. Nat Neurosci 11, 505–513 (2008). 10.1038/nn2070

60 Shibata, K., Sasaki, Y., Kawato, M. & Watanabe, T. Neuroimaging Evidence for 2 Types of Plasticity in Association with Visual Perceptual Learning. Cereb Cortex 26, 3681–3689 (2016). 10.1093/cercor/bhw176

61 Liu, J., Dosher, B. A. & Lu, Z. L. Augmented Hebbian reweighting accounts for accuracy and induced bias in perceptual learning with reverse feedback. J Vis 15, 10 (2015). 10.1167/15.10.10

62 Jenkinson, M., Bannister, P., Brady, M. & Smith, S. Improved optimization for the robust and accurate linear registration and motion correction of brain images. Neuroimage 17, 825–841 (2002).

63 Jenkinson, M. & Smith, S. A global optimisation method for robust affine registration of brain images. Med Image Anal 5, 143–156 (2001).

64 Andersson, J. L., Jenkinson, M. & Smith, S. Non-linear registration, aka Spatial normalisation FMRIB technical report TR07JA2. FMRIB Analysis Group of the University of Oxford 2, 1–21 (2007).

65 Andersson, J. L., Jenkinson, M. & Smith, S. Non-linear optimisation. FMRIB technical report TR07JA1. Practice. 2007a Jun (2007).

66 Jenkinson, M., Bannister, P., Brady, M. & Smith, S. Improved optimization for the robust and accurate linear registration and motion correction of brain images. Neuroimage 17, 825–841 (2002).

67 Boynton, G. M., Engel, S. A., Glover, G. H. & Heeger, D. J. Linear systems analysis of functional magnetic resonance imaging in human V1. J Neurosci 16, 4207–4221 (1996). 10.1523/JNEUROSCI.16-13−04207.1996

68 Heeger, D. J., Huk, A. C., Geisler, W. S. & Albrecht, D. G. Spikes versus BOLD: what does neuroimaging tell us about neuronal activity? Nat Neurosci 3, 631–633 (2000). 10.1038/76572

69 Ogawa, S. et al. Intrinsic signal changes accompanying sensory stimulation: functional brain mapping with magnetic resonance imaging. Proc Natl Acad Sci U S A 89, 5951–5955 (1992). 10.1073/pnas.89.13.5951

70 Fischl, B. et al. Automatically parcellating the human cerebral cortex. Cerebral Cortex 14, 11–22 (2004). 10.1093/cercor/bhg087

71 Glasser, M. F. et al. A multi-modal parcellation of human cerebral cortex. Nature 536, 171–178 (2016). 10.1038/nature18933

72 Friston, K. J. et al. Analysis of fMRI time-series revisited. Neuroimage 2, 45–53 (1995). 10.1006/nimg.1995.1007

73 Worsley, K. J. & Friston, K. J. Analysis of fMRI time-series revisited--again. Neuroimage 2, 173–181 (1995). 10.1006/nimg.1995.1023

74 JASP (Version 0.17.3)[Computer software] (2023).

75 Morey, R. D. Confidence intervals from normalized data: A correction to Cousineau (2005). Tutorials in Quantitative Methods for Psychology 4, 61–64 (2008).

